# Integrated Bioinformatics Analysis Reveals Candidate Hub Genes and Regulatory Networks Associated with Preeclampsia

**DOI:** 10.64898/2026.07.20.739688

**Authors:** Dan Jiang, Haiyan Xiong, Qingqing Li, Zixin Pi

**Author notes:** Correspondence:. Department of Obstetrics and Gynecology, The First People’s Hospital of Jiangxia District, Wuhan, Hubei University of Medicine. No. 1, Wenhua Avenue, Jiangxia District, Wuhan 430200, Hubei Province, China. These authors contributed equally to this work (Dan Jiang, Haiyan Xiong).

## Abstract

Preeclampsia (PE) is a severe pregnancy-associated hypertensive disorder and a major contributor to maternal and perinatal morbidity and mortality. The mechanisms of PE pathogenesis are not yet understood. This paper aims to explore candidate PE-associated biomarkers and regulatory mechanisms using bioinformatics analysis of placental transcriptomic datasets. We downloaded placental transcriptomic datasetsGSE203507 and GSE148241 from GEO and investigated differentially expressed genes(DEGs) between PE and control samples. We performed Gene Ontology (GO), Kyoto Encyclopedia of Genes and Genomes (KEGG), Disease Ontology (DO), and protein–protein interaction (PPI) analyses to characterize the potential role, disease associations, and interaction networks of the DEGs. We validated the mRNA expression patterns of candidate hub genes using two independent placental transcriptomic datasets, GSE143966 and GSE114691.We identified 263 DEGs in the discovery analysis and further obtained 150 overlapping DEGs including80 upregulated genes and 70 downregulated genes, for downstream analysis. We then identified eight candidate hub genes: OPRK1, OXGR1, HCAR3, CCR5, HCAR2, CXCL1, FPR3, and SSTR1using Metascape software(v3.5.20260201). In the validation phase, most candidate hub genes showed broadly consistent mRNA expression trends across GSE114691 and GSE143966, while FPR3 showed weaker cross-dataset consistency. These findings provide candidate PE-associated genes and regulatory pathways for further experimental and clinical validation..

## Introduction

Preeclampsia (PE) is a pregnancy-specific multisystem disorder defined by the onset of hypertension and proteinuria occurring after 20 weeks of gestation. It affects approximately 2% to 8% of pregnancies globally and represents a major cause of maternal and perinatal mortality [1].

Although the cause of preeclampsia is still debated, clinical and pathological studies indicate that the placenta plays an important role in the pathogenesis of the syndrome [2]. Placental dysfunction is widely considered a central feature of PE, involving impaired spiral artery remodeling, abnormal trophoblast invasion, placental ischemia/hypoxia, dysregulated maternal inflammatory responses, and endothelial dysfunction[3]. These interconnected processes indicate that PE is unlikely to be driven by a single molecular event. Instead, integrated transcriptomic analysis may help identify changes in coordinated gene-expression and regulatory pathways that lead to placental dysfunction and PE development. The rapid development of placental tissue sequencing technology has promoted the identification of candidate biomarkers and molecular pathways potentially associated with PE [4,5]. Bioinformatics analyses of publicly available transcriptomic datasets provide new insights to identify disease-associated genes, transcripts, and regulatory pathways. Reanalysis of data sets from independent cohorts may help clarify the molecular networks associated with PE. Recent studies of PE using RNA sequencing and bioinformatics tools have revealed dysregulated pathways such as the G protein-coupled receptor (GPCR) signaling pathway, endocytosis pathway, focal adhesion pathway, and immune/inflammatory pathways, alongside abnormal expression of various miRNAs and mRNAs [6–8]. These discoveries have greatly improved our understanding of the pathophysiology of PE. identifying robust PE-associated molecular markers remains challenging because many studies are heterogeneous datasets, limited by small sample sizes, insufficient cross-dataset validation, and incomplete clinical subtype information. Therefore, this study integrated multiple placental publicly available transcriptomic datasets to identify candidate PE-associated hub genes and regulatory networks and to assess their expression patterns in independent validation datasets.

## 2. Results

### 2.1. Identification of DEGs in PE

We analyzed GEO placental transcriptomic datasets to determine the differentially expressed genes (DEGs) between PE and control pregnancies. A flowchart outlining the screening approach for identifying these genes is presented in Figure 1. Using the criteria described in the Materials and Methods section, DEGs were identified and illustrated using volcano plots for each discovery dataset(Figure 2A,B; Supplementary Figure 1). The analysis revealed 263 DEGsin the discovery datasets; among them, 150 overlapping DEGs, including 80 upregulated and 70 downregulated genes, were retained for downstream analysis, as depicted in Figure 2C,D.

**Figure 1.**
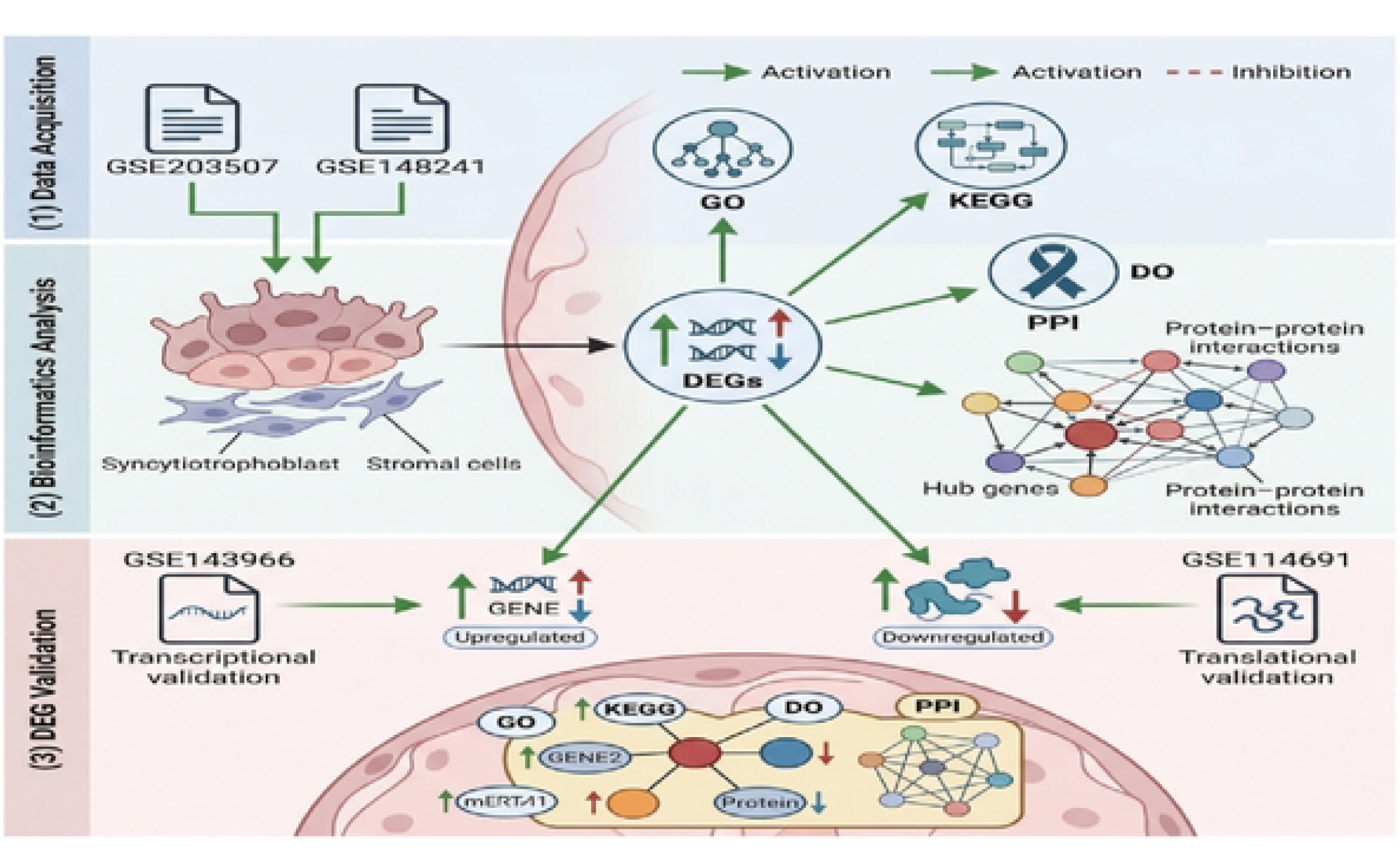
Flowchart of the overall bioinformatic analysis

**Figure 2.**
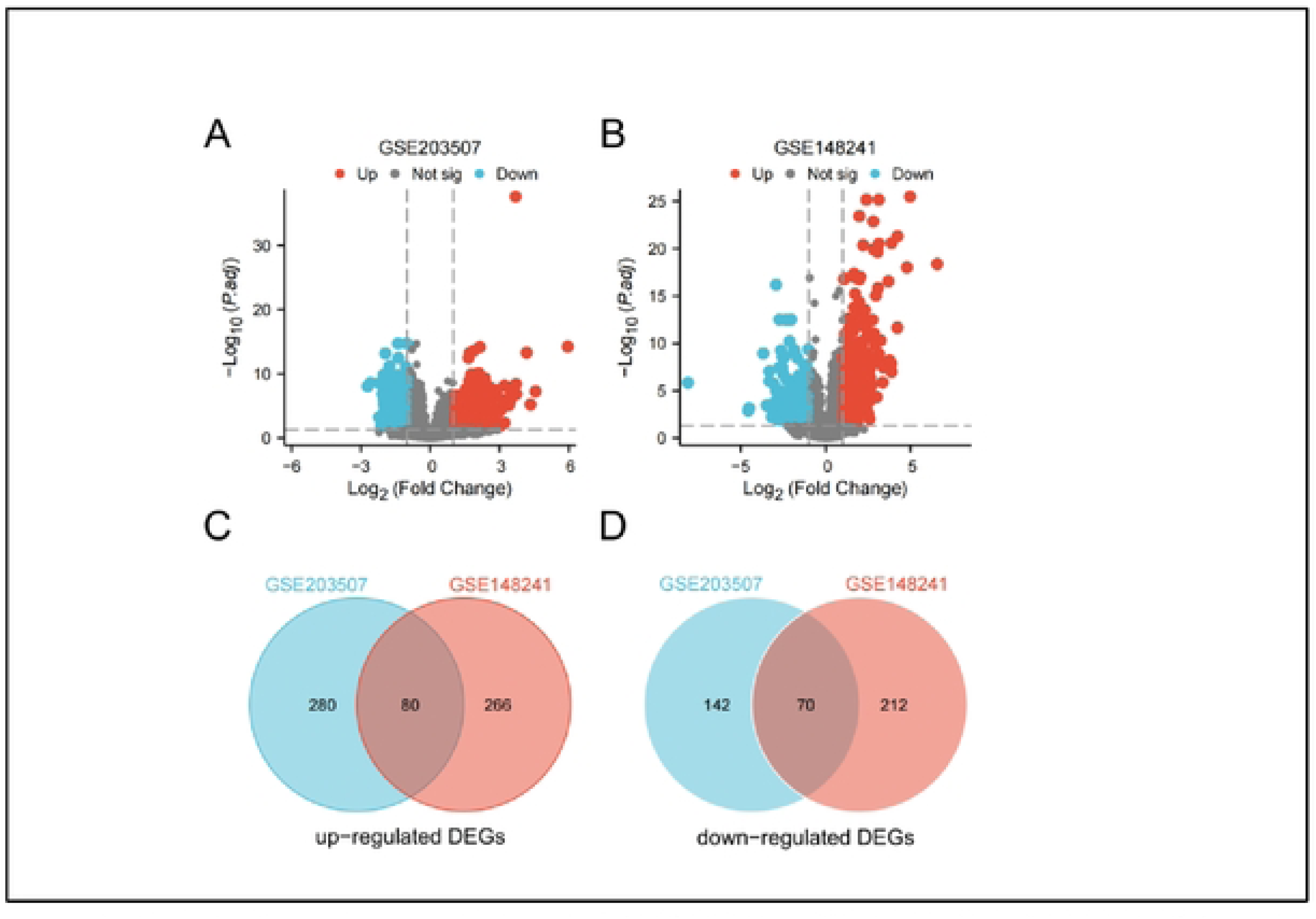
Identification of common DEGs in PE. (A,B) Volcano plots of DEGs in GSE203507 (A) and GSE148241 (B). (C,D) Venn diagrams of overlapping upregulated (C) and downregulated (D) DEGs.

### 2.2. GO and KEGG Enrichment Results of DEGs

We characterized the functional and regulatory networks of differentially expressed genes (DEGs) in PE based on multi-dimensional enrichment analysis. DEGs were mainly enriched in pathways related to hypoxia response (HIF-1 pathway), cell adhesion, MAPK signaling, reproductive endocrine regulation, and immune homeostasis, suggesting that multiple biological processes may be involved in PE-related placental dysfunction (Figure 3A,B). Disease Ontology enrichment analysis showed association with preeclampsia-related terms, such as hypoxia, pregnancy-induced hypertension, and preeclampsia, as well as pregnancy-related outcomes such as miscarriage and endocrine disorders (Figure 3D). Cell-type enrichment analysis revealed that DEGs were predominantly expressed in placental trophoblasts and hepatic cells-related signatures, which may reflect links between placental dysfunction, metabolic regulation, and inflammatory processes(Figure 3C). Transcription factor target enrichment identified key regulatory nodes like HIF-1, RBL1, and CEBPA, suggesting that aberrant upstream transcriptional regulation may contribute to DEG expression in PE (Figure 3E). Together, these results summarize the molecular pathological landscape of PE at functional, phenotypic, cellular, and regulatory levels, allowing us to understand its pathology and explore candidate biomarkers and molecular mechanisms.

**Figure 3.**
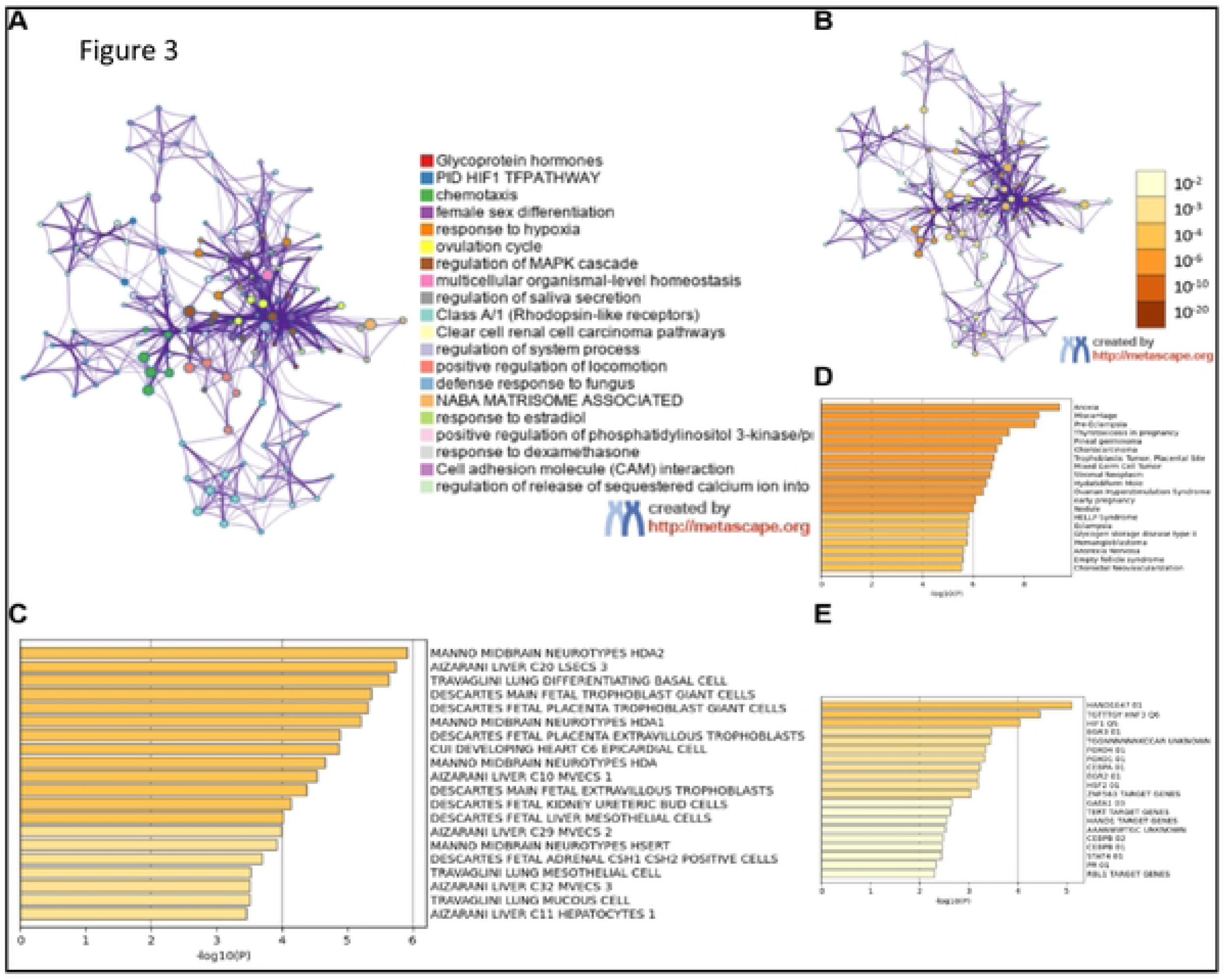
Functional enrichment and regulatory analysis of DEGs. (A) PPI network colored by functional pathways. (B) Disease-associated PPI network. (C) Cell-type enrichment. (D) Disease ontology enrichment. (E) Transcription factor target enrichment.

### 2.3. Protein–Protein Interaction (PPI) Network Construction

A PPI network was constructed using the candidate GED set, resulting in a network consisting of 75 nodes and 482 edges with scale-free topological features. Highly connected hub genes, namely, OXGR1, HCAR3, CXCL1, and CCR5, were identified as candidate hub genes in the PPI network (Figure 4A). MCODE module analysis identified three densely connected subnetworks with potential functional relevance: Module 1 consisted of eightgenes,OXGR1, OPRK1, HCAR3, FPR3, CCR5, CXCL1, SSTR1, HCAR2,in a densely connected cluster for inflammatory and immune signaling (Figure 4B); Module 2 consists of four genes (CGA, CGB3, CGB5, CGB8), forming a compact subnetwork for glycoprotein hormone signaling (Figure 4C); Module 3 consisted of threegenes,CLDN16, CLDN1, CLDN10, forming a tight junction-related modules for epithelial barrier integrity (Figure 4D).

**Figure 4.**
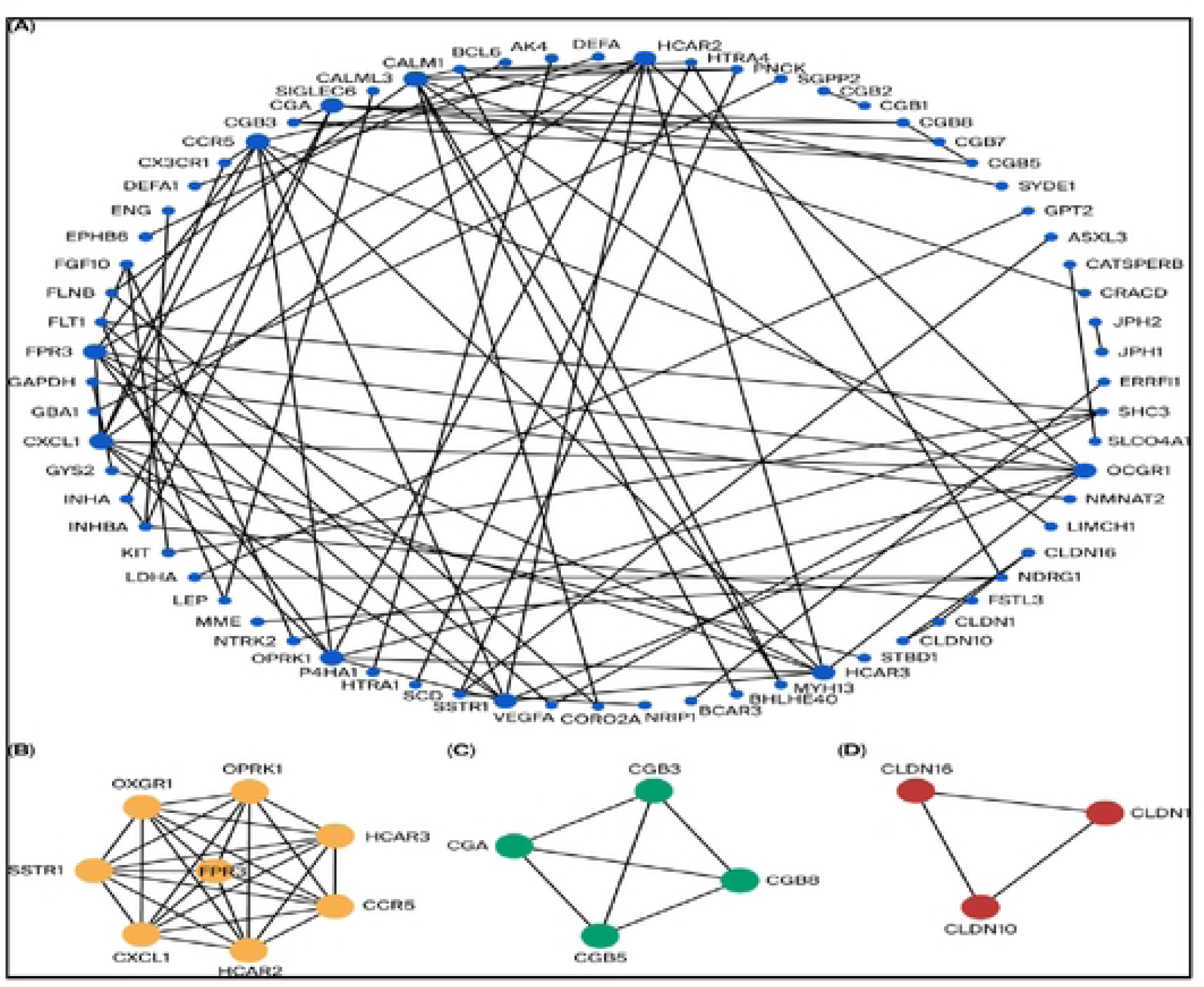
PPI network and module analysis. (A) Global PPI network. (B) Core hub gene subnetwork. (C) Glycoprotein hormone subnetwork. (D) Tight junction subnetwork.

### 2.4. Dataset Validation of 8 Hub Genes’ Expression

To explore the expression patterns and diagnostic value of eight candidate genes (OPRK1, OXGR1, HCAR3, CCR5, HCAR2, CXCL1, FPR3, SSTR1) in this disease, we compared the PE group with the control group and analyzed their expression patterns and diagnostic discrimination in GSE143966 and GSE114691 validation datasets. In GSE143966, the expression of OXGR1, HCAR3, CCR5, HCAR2, CXCL1, and FPR3 in the PE group was significantly upregulated (*p*< 0.05), while the expression of OPRK1 and SSTR1 was downregulated (*p*< 0.05) (Figure 5F). In GSE114691, the expression of OXGR1, HCAR3, CCR5, HCAR2, and CXCL1 was also significantly upregulated (*p*< 0.05), and the expression of OPRK1 and SSTR1 was downregulated (*p*< 0.05), which was consistent with the trend observed in GSE143966. In contrast, FPR3 showed an inconsistent expression trend inGSE114691 compared withGSE143966 (Figure 5C), suggesting weaker cross-dataset reproducibility. However, the relatively small sample size of GSE143966 should be considered when interpreting the expression and ROC results. The diagnostic value was further evaluated via the receiver operating characteristic (ROC) curve. In GSE114691 (Figure 5A,B), HCAR2 demonstrated a stable and favorable diagnostic discrimination in GSE114691 (AUC = 0.785). HCAR3 (AUC = 0.718) and CXCL1 (AUC = 0.710) showed moderate-to-good dignostic discrimination, while FPR3 (AUC = 0.523) demonstrated limited independent diagnostic value. Among GSE143966 (Figure 5D,E), OPRK1 has the highest diagnostic efficacy (AUC = 0.906), followed by CCR5 (AUC = 0.852), HCAR2 (AUC = 0.812), and OXGR1 (AUC = 0.781). Cross-dataset analysis revealed that all seven genes had some diagnostic efficacy (AUC > 0.6) except FPR3. The OPRK1 value was relatively high (AUC of GSE143966 = 0.906), meaning that it might be the best diagnostic marker.

**Figure 5.**
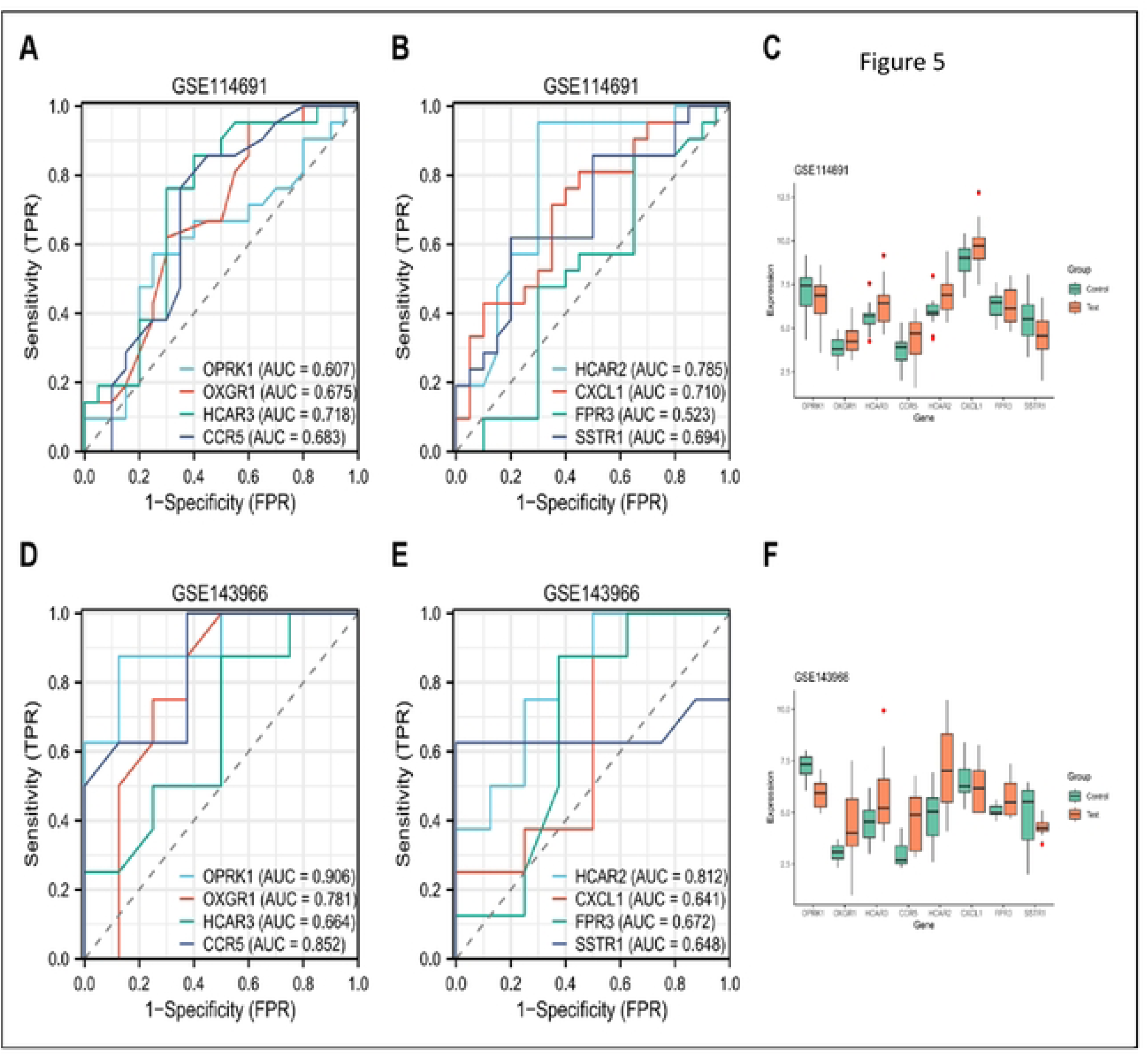
Expression and diagnostic performance of hub genes. (A,B) ROC curves in GSE114691. (C) Expression boxplots in GSE114691. (D,E) ROC curves in GSE143966. (F) Expression boxplots in GSE143966

### 2.5. miRNA–Hub Gene and TF–Hub Gene Networks

In this study, we constructed a multi-level regulatory network comprising microRNAs (miRNAs), transcription factors (TFs), and eight hub genes in preeclampsia (PE) (Figure 6A). At the transcriptional level, we identified 14 key TFs: CXCL1 was predicted to interact with eight TFs; CCR5 was predicted to interact with five TFs; and SSTR1 was linked only to EZH2 in the TF-hub gene network (Figure 6B). At the post-transcriptional level, we identified 227 miRNAs targeting these hub genes. Notably, SSTR1 had the greatest number of miRNAs (65). CXCL1, OPRK1, and other hub genes were predicted to be targeted by 58, 42, and 8-17 miRNAs, respectively (Figure 6C).

**Figure 6.**
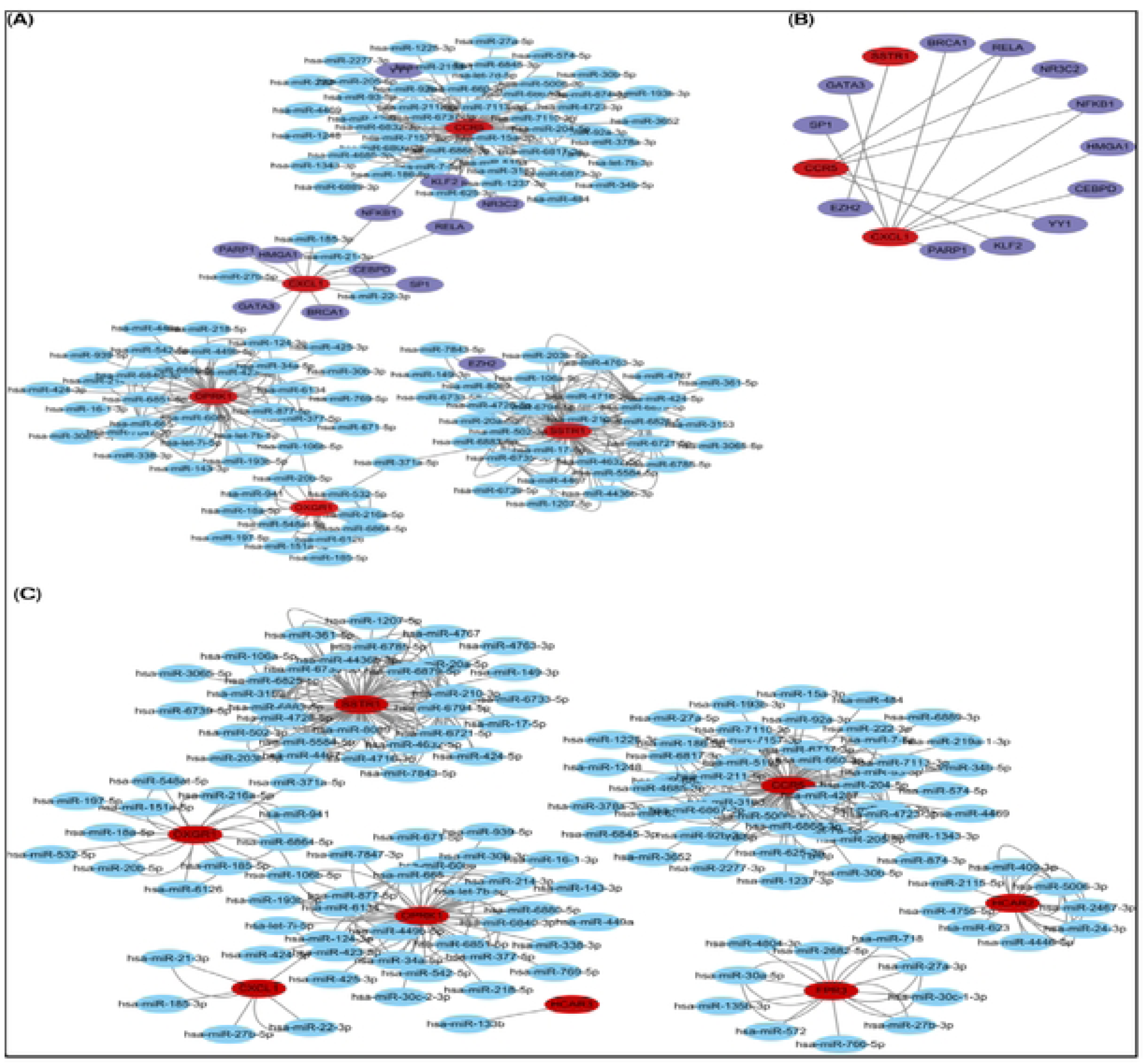
Regulatory network analysis of hub genes. (A) Global miRNA - target gene - TF network. (B) TF - core gene subnetwork. (C) Focused miRNA - core gene subnetwork.

## 3. Discussion

PE is a pregnancy-specific hypertensive disorder and a major cause of complications and mortality during gestation [26]. Around 76,000 women die annually from PE, representing 16% of maternal deaths, with the highest burden being in developing nations [27]. However, the molecular mechanisms involved in PE are incompletely understood, and it is difficult to advance early prediction, diagnosis, and therapy. Identifying key biomarkers is essential for the identification, diagnosis, and management of PE. This study combined two GEO datasets related to PE for bioinformatics analysis to explore potential signaling pathways and biomarkers. In addition, we used another two GEO datasets associated with PE to validate the biomarkers.

The Disease Ontology (DO) analysis showed that the DEGs were enriched in PE-related disease terms. This suggests that the DEGs identified in this study are closely related to PE and are expressed in severe conditions such as eclampsia and HELLP syndrome. The functional enrichment analysis uncovered important biological processes and signaling pathways involved in the molecular pathogenesis of preeclampsia. The enrichment of the PID HIF1 TF pathway (Log10(P) = −9.92) and the GO term “response to hypoxia” (Log10(P) = −6.74) were the most significantly enriched signatures, strongly establishing placental ischemia and hypoxia-driven HIF1 signaling as central initiators of PE. These pathways also fuel downstream inflammatory chemotaxis (Log10(P) = −8.92) and PI3K/Akt signaling activation, which, together, promote endothelial dysfunction and aberrant immune cell invasion, hallmarks of PE-related systemic vascular injury [28–30]. The enrichment of the “female differentiation,” “response to estradiol,” and “ovulation cycle” highlight that hormonal dysregulation plays a key role in compromising placental development and vascular homeostasis. The NABA matrisome-associated pathway and KEGG pathway’s “cell adhesion molecule (CAM) interaction” points to impaired extracellular matrix (ECM) remodeling and cell adhesion as critical mechanisms driving shallow trophoblast invasion and resultant placental insufficiency. Moreover, enrichment of the Class A/1 (rhodopsin-like receptors) Reactome gene sets resembles our identified hub genes (e.g., OPRK1, OXGR1, SSTR1), linking dysregulated G protein-coupled receptor (GPCR) signaling to vasomotor dysfunction, inflammation, and hormonal imbalance in PE. This is consistent with previous studies in preeclampsia [31–33]. Together, these findings suggest thathypoxia, inflammation, hormonal dysregulation, and ECM remodeling, offering mechanistic insights into PE pathophysiology and nominating promising candidates for future diagnostic and therapeutic exploration.

This study identified eight candidate hub genes, includingOPRK1, OXGR1, HCAR3, CCR5, HCAR2, CXCL1, FPR3, SSTR1, based on PPI and module analyses. The validated data showed that the mRNA expression of OPRK1, OXGR1, HCAR3, CCR5, HCAR2, CXCL1, and SSTR1 genes differs between PE and healthy maternal placenta. Previous studies have reported that OPRK1 [34], CXCL1 [35], and CCR5 [36,37] are biomarkers of PE. OXGR1, an intermediate in GPCRs, plays a role in the citric acid cycle as a receptor for α-ketoglutarate and metabolic processes. GPCRs can regulate cell proliferation by modulating the downstream signaling of G protein signaling pathways. HCAR3, classified as a class A G-protein-coupled receptor, functions as a sensor of cell energy metabolism and plays a significant role in the regulation of lipolysis in humans. This receptor plays a role in various physiological processes and is a promising target for the treatment of metabolic disorders, tumors, and immune diseases [38]. HCAR2 mediates different pathophysiological events by activating G protein or β-arrestin effectors, influenced by endogenous ketone body β-hydroxybutyrate and exogenous niacin, and is a promising therapeutic target for inflammation-related diseases [39]. Accumulating evidence indicates that inflammation-related diseases play a significant role in the pathogenesis of preeclampsia [40–42]. β-arrestin effectors refer to the downstream signaling molecules, kinases, scaffold proteins, and endocytic and ubiquitination machinery recruited, bound, and regulated by β-arrestins (β-arrestin 1/2). As executors of GPCR signaling from G protein, they are dependent on β-arrestin-dependent pathways [43]. SSTR1 is a member of the G protein-coupled receptor family. Its functions include inhibiting hormone secretion and regulating cell proliferation and apoptosis, and it participates in neural, endocrine, and tumor regulation [44]. It has been reported that SSTR1 can be used as a predictor of obesity [45,46]. Obesity has been shown to be associated with preeclampsia, and the effect of obesity on preeclampsia varies with its severity [47–49]. In summary, OXGR1, HCAR3, HCAR2, and SSTR1 facilitate pathophysiology via signaling pathways. In recent years, new evidence has indicated that the G-protein signaling pathway plays a role in preeclampsia [50,51]. Based on this analysis, OXGR1, HCAR3, HCAR2, and SSTR1 are biomarkers of PE, particularly HCAR2, HCAR3, and OXGR1, which have an area under the ROC curve (AUC) of up to 70%, suggesting potential diagnostic discrimination in these datasets, although larger independent cohorts are required to confirm their diagnostic utility.

To further investigate the regulatory mechanisms of the candidate hub genesin PE, we constructed predicted TF-hub gene and miRNA–hub gene regulatory networks (Figure 6A), identifying 14 key transcription factors (TFs) (Figure 6B) and 227 microRNAs (miRNAs). SSTR1 has the greatest number of interacting miRNAs, indicating that its post-transcriptional fine-tuning process is complex. MiRNAs targeting CXCL1, OPRK1, and other hub genes have redundant and coordinated regulatory patterns (Figure 6C). We found that NFKB1, RELA, and hsa-miR-371a-5p in the transcription factor target genes were closely associated with the hub genes, which is consistent with previous related reports [52–54]. Comprehensive analysis reveals that this regulatory network has established a core signaling module centered on inflammation, immune infiltration, and vascular dysfunction. Core TFS, such as RELA and NFKB1, work in synergy with miRNA to regulate key hub genes, such as CXCL1 and CCR5. This provides a theoretical basis for developing a deeper understanding of the molecular pathogenesis of PE at the systematic level and for exploring potential diagnostic and therapeutic targets.

Several limitations should be acknowledged. First, detailed clinical information, including PE subtype, gestational age, maternal BMI, fetal growth restriction status, medication use, and other potential confounders, was not uniformly available across datasets. Second, this study was based on retrospective analysis of public transcriptomic datasets with heterogeneous platforms and relatively small sample sizes. Third, the findings were validated only at the mRNA level, without functional experiments or protein-level validation. Therefore, the candidate hub genes and predicted regulatory networks require further confirmation using independent clinical cohorts, qRT-PCR, protein-level assays, and functional studies in relevant placental cell models.

## 4. Materials and Methods

### 4.1. Expression Profile Dataset Selection

Placental transcriptomic datasets related to preeclampsia were obtained from the GEO database (https://www.ncbi.nlm.nih.gov/geo/). Datasets were included if they contained human placental transcriptomic profiles from PE and control pregnancies with available case-control grouping information. Datasets were excluded if they were not based on human placental tissue, did not provide accessible expression data for reanalysis, or lacked of appropriate control samples. The GSE203507 series [9], including data on 16 preeclampsia and 10 uncomplicated term births, and the GSE148241 dataset [10], including data on 9 PE and 34 uncomplicated term births, were chosen as discovery datasets. Two datasets, namely, GSE114691 [11], including data on 20 PE and 21 uncomplicated term births, and GSE143966 [12], including data on 8 PE and 8 uncomplicated term births, were included in the validation sets analysis (Table 1).Where available, dataset-level information, including platform, sample size, grouping, and tissue source, was extracted. Detailed clinical variables such as gestational age, fetal growth restriction status, PE subtype, maternal BMI, ethnicity, parity, and medication use were not uniformly available across all datasets and were therefore considered as potential limitations.

**Table 1.** The Summary of Datasets Used in This Study Summary of the four GEO datasets used in this study, including platform, tissue type,experiment type, sample size, and grouping.

| GEO Accession | Platform | Tissue Type | Experiment type | Samples | Attribute |
| --- | --- | --- | --- | --- | --- |
| GSE203507 | GPL16791 | Human placenta | high throughput sequencing | 16vs10 | Test set |
| GSE148241 | GPL16791 | Human placenta | high throughput sequencing | 9vs34 | Test set |
| GSE114691 | GPL11154 | Human placenta | High throughput sequencing | 20vs21 | Validation set |
| GSE143966 | GPL20301 | Human placenta | high throughput sequencing | 8vs8 | Validation set |

### 4.2. Data Preprocessing

Series matrix files were downloaded and processed using R version 4.2.2.When required, expression values were log2-transformed before differential expression analysis. Gene identifiers were mapped to official gene symbols using the corresponding platform annotation files and Bioconductor packages. For genes represented by multiple probes or entries, the mean expression value was used. DEGs were identified using the limma package inR/Bioconductor.Genes with |log2FC| > 1 and an adjusted p value/FDR < 0.05 were considered significant DEGs. Volcano plots were created using GraphPad prism version 10.6.1 (GraphPad Software, San Diego, CA, USA). The common up- and downregulated significant DEGs shared between the GSE203507 and GSE148241 datasets were overlapped with an online Venn tool (http://www.bioinformatics.com.cn/static/others/jvenn/example.html). Enrichment analyses via Gene Ontology (GO) and the Kyoto Encyclopedia of Genes and Genomes (KEGG) pathway of DEGs were conducted via Meta scape (Meta scape Gene List Analysis Report V3.5.20260201), which has contributed extensively to our comprehension of gene functional categories.

### 4.3. Pathway and Process Enrichment Analysis

Pathway and process enrichment were performed using several databases: KEGG Pathway, Gene Ontology Biological Processes, Reactome Gene Sets, Canonical Pathways, CORUM, WikiPathways, and PANTHER Pathway. Allgenes in the Meta scape database were used as the background set(Metascape v3.5.20260201). Enrichment *p*-values were computed based on the cumulative hypergeometric distribution [13], and q-values were obtained using the Benjamini– Hochberg procedure to adjust for multiple testing [14]. Terms were considered significantly enriched according to the Metascape default criteria of *p*-value < 0.01, minimum gene count o≥ 3, and enrichment factor > 1.5, defined as the ratio of observed counts to expected counts. To minimize redundancy, enriched terms were clustered according to gene membership similarities, employing the kappa coefficient [15] as the similarity metric for hierarchical clustering. Subtrees with a similarity score > 0.3 were classified as clusters, and the most statistically significant term within each cluster was designated as the cluster representative. The resulting network of enriched terms was visualized using Metascape [16] (v3.5.20260201) by cluster ID and *p*-value. Additionally, gene list enrichments were identified in the following ontology categories: Cell_Type_Signatures, DisGeNET, and Transcription_Factor_Targets.

### 4.4. Protein–Protein Interaction Enrichment Analysis

Protein–protein interaction (PPI) networks were used to explore potential physical or functional associations among proteins encoded by the candidate genes at the protein level, enhancing comprehension of disease mechanisms, and assisting in target identification. Protein– protein interaction enrichment analysis was performed using databases such as STRING [17], BioGrid [18], OmniPath [19], and InWeb_InBioMap2 [20]. Specifically, interactions with a score exceeding 0.132 in STRING and those verified in BioGrid are considered. However, this relatively low threshold may include weak functional associations and was considered a methodological limitation. The PPI network is then visualized using Metascape software.

### 4.5. Identifying the Hub Genes and Key Module

Candidates hub genes were identified from the PPI network through Meta scape and cytoHubba, and Molecular Complex Detection (MCODE) was used to identify densely connected modules. The Cytoscape plugin CytoHubba was also used to find hub genes in the PPI network using five topological algorithms: Degree, MNC (Maximum Neighborhood Component), DMNC (Density of Maximum Neighborhood Component), MCC (Maximal Clique Centrality), and EPC (Edge Percolated Component). The top 10 genes ranked by each cytoHubba algorithm were recorded, and genes consistently ranked across multiple algorithms and/or located in the core MCODE module were selected as final candidate hub genes, while the following parameters were used: node score cutoff = 0.2, degree cutoff = 2, K-core = 4, and max depth = 100 (Table 2).

**Table 2.**
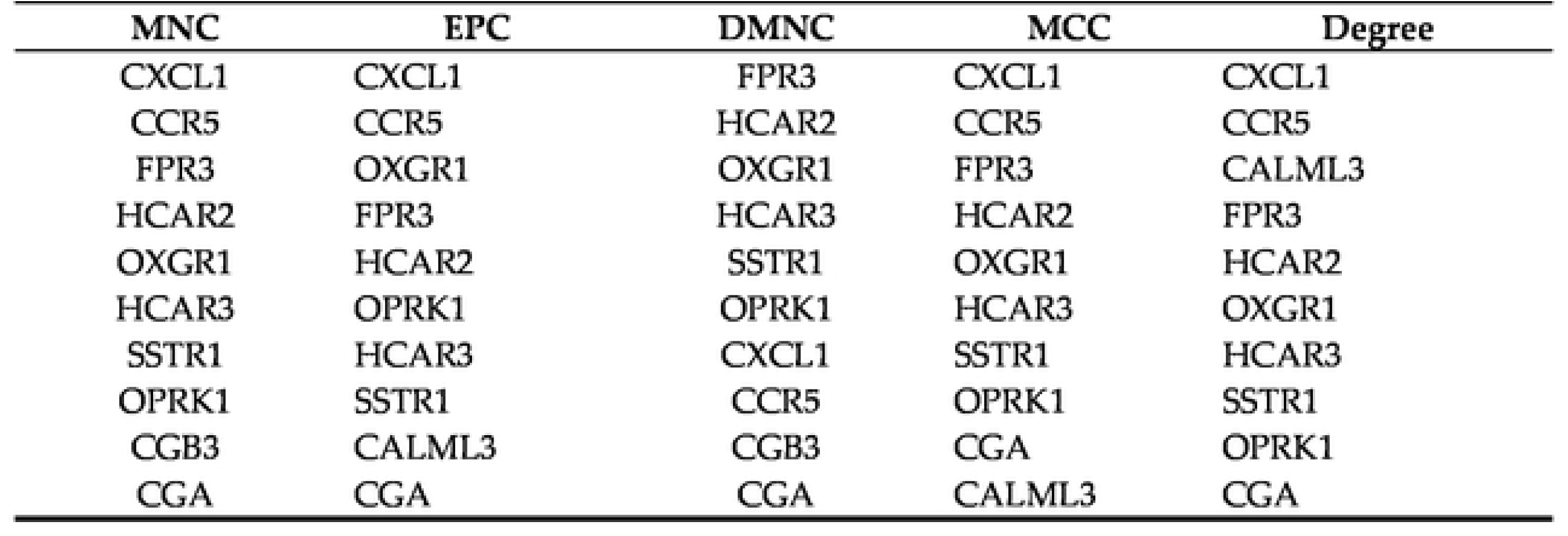
The Top 10 Hub Genes Analyzed by cyto Hubba Top 10 hub genes identified by topological algorithms (MNC, EPC, DMNC, MCC, Degree) in cytoHubba.

| MNC | EPC | DMNC | MCC | Degree |
| --- | --- | --- | --- | --- |
| CXCL1 | CXCL1 | FPR3 | CXCL1 | CXCL1 |
| CCR5 | CCR5 | HCAR2 | CCR5 | CCR5 |
| FPR3 | OXGR1 | OXGR1 | FPR3 | CALML3 |
| HCAR2 | FPR3 | HCAR3 | HCAR2 | FPR3 |
| OXGR1 | HCAR2 | SSTR1 | OXGR1 | HCAR2 |
| HCAR3 | OPRK1 | OPRK1 | HCAR3 | OXGR1 |
| SSTR1 | HCAR3 | CXCL1 | SSTR1 | HCAR3 |
| OPRK1 | SSTR1 | CCR5 | OPRK1 | SSTR1 |
| CGB3 | CALML3 | CGB3 | CGA | OPRK1 |
| CGA | CGA | CGA | CALML3 | CGA |

### 4.6. Verification of Hub Gene Expression by Validation Datasets

The validation datasets GSE114691 and GSE143966 were processed using the same annotation, transformation, and normalization procedures applied to the discovery datasets where applicable. The mRNA expression level of the candidate hub genes were compared between PE and control samples in GSE114691 and GSE143966. ROC curve analysis was carried out to assess the diagnostic discrimination of each candidate hub gene, and the area under the curve (AUC) was calculated.

### 4.7. Construction of miRNA–Hub Gene and TF–Hub Gene Networks

Target miRNAs of the hub genes were predicted using the miRTarBase [21] (v 9.0), Starbase [22] v9.0, and miWalk [23] v3.0 databases. To enhance prediction accuracy, we selected miRNAs for each hub gene that were identified by at least two of the databases. MiRNAs that regulate multiple hub genes were considered potentially important regulatory miRNAs. Candidate hub genes were queried in TRRUST [24] v2 platform for transcription factor (TF) prediction. The miRNAs-hub gene and TFs-hub gene networks were visualized using Cytoscape [25] v3.10.4.

### 4.8. Statistical Analysis

Statistical analyses were performed using R softwareR4.2.2) and GraphPad Prism(10.6.1). The R package limma was employed to detect DEGs in RNA-seq datasets. Differential expression analysis, validation comparisons, and ROC analyses were performed as described above. Unless otherwise specified, p < 0.05 was considered statistically significant. For genome-wide DEG screening, adjusted p value/FDR correction should be applied where available.

## 5. Conclusions

The findings of this study provide additional insight additional insight into PE-related placental transcriptomic changes and suggest candidate genes and regulatory pathways for further investigation. The identified genes, particularly OXGR1, HCAR3, HCAR2, and SSTR1, may represent candidate PE-associated genes, but their diagnostic and therapeutic relevance requires further validation. Future studies should validate these genes in independent placental cohorts, assess protein-level expression using immunohistochemistry, Western blotting, or ELISA, and investigate their biological roles in placental cell models under hypoxic or inflammatory conditions.

## Author Contributions

Conceptualization, A Methodology, Software, Formal Analysis, Visualization, Resources, Data Curation, Writing and-Original Draft, ZixinPI; Writing, Review and Editing, DanJiang; Supervision, Haiyan Xiong; Validation, Qingqing Li.

## Funding

No funding was received for this work

## Data Availability Statement

All datasets analyzed in this study are publicly available in the GEO database (https://www.ncbi.nlm.nih.gov/geo/), with accession numbers GSE203507, GSE148241, GSE114691, and GSE143966.

## Ethical Statement

Ethical approval was not required because all data were obtained from public transcriptome databases.

## Conflicts of Interest

The authors declare no conflict of interest.

We confirm that neither the manuscript nor any parts of its content are currently under consideration for publication with or published in another journal.

All authors have approved the manuscript and agree with its submission to PLOS ONE.

## Notes

### Competing Interest Statement

The authors have declared no competing interest.

